# Drebrin-like protein regulates body bending of *C. elegans* via suppression of NCA cation leak channels

**DOI:** 10.1101/2019.12.12.874289

**Authors:** Eugenia Butkevich, Peter Weist, Daniel Härtter, Dieter R. Klopfenstein, Renata Garces, Christoph F. Schmidt

## Abstract

Drebrin-like protein (DBN-1) in *C. elegans* is an adaptor protein that connects different cellular pathways to the actin cytoskeleton. Using a CRISPR-Cas9 system, we generated a new *dbn-1* allele, which lacks 80% of C-terminal part of DBN-1. The mutant displays a striking hyper-bending locomotion phenotype and body posture with two times stronger curvature than wild type. We show by atomic force microscopy that the muscle tone of the mutant remains unaffected. Aiming to track down the cause of hyper-bending, we performed genetic epistasis experiments. We found that mutations in the Rho-specific guanine-nucleotide exchange factor (GEF) domain of UNC-73 (Trio), pan-neuronal expression of dominant negative RHO-1 and mutations in NCA (NALCN) cation leak channels all suppressed hyper-bending in the *dbn-1* mutant. These data indicate that DBN-1 negatively regulates the activity of both NCA-1 and NCA-2 channels, opposing RHO-1 in the non-canonical Gq pathway. We conclude that DBN-1 is an important component of the neuronal signaling cascade that controls the degree of *C. elegans* body bending during locomotion.

## Introduction

*C. elegans* nematodes move over firm substrates lying on their side and bending their body in sinusoidal waves ^1, 2^. The bending results from alternating contractions and relaxations of ventral and dorsal groups of body-wall muscles ^3^. Muscle contraction is controlled by the activity of excitatory and inhibitory motor neurons: excitation of cholinergic neurons induces muscles contraction at one side of the body while at the same time, these neurons excite inhibitory GABA neurons relaxing muscles on the opposite side of the body ^4,5^. The force produced by body-wall muscles is transmitted via dense bodies to the underlying elastic cuticle driving the movement of the entire animal ^6^.

One of the central roles in the regulation of neuronal activity that controls *C. elegans* locomotion is played by NCA sodium leak channels. NCA/NALCN channels are nonselective, voltage independent, and do not inactivate. Providing the dominant basal sodium leak conductance, they set the resting membrane potential and neuronal excitability ^7^. The NCA conductance mediates propagation of neuronal activity from the cell bodies to synapses and synaptic transmission at neuromuscular junctions required for normal locomotion ^8, 9^. The NLF-1 (NCA localization factor-1) protein delivers NCA channels to the plasma membrane ^10^ where several signal transduction pathways regulate their activity. The M3 muscarinic receptor and the TACR1 (G-protein-coupled receptor for substance P) both activate conductance through NALCN channels via the Src tyrosine kinase-dependent pathway ^11, 12^. Another set of GPCRs, the CaSR (calcium-sensing receptor), GABA-BR (gamma-aminobutyric acid-B receptor) and D2R (dopamine receptor) inhibits NALCN activity ^13, 14^. In addition, NALCN modulates neuronal membrane potential in co-operation with potassium channels and gap junction hemichannels ^15, 16^. Recently, a new pathway regulating NCA channels in *C. elegans* was discovered ^17, 18, 19^. It was found that the heteromeric Gq protein activates UNC-73/ RhoGEF Trio, which subsequently activates the small GTPase RHO-1. RHO-1 then acts via an unknown mechanism to activate the NCA ion channels ^17^.

Drebrin-like protein (known as mAbp1, HIP-55, SH3P7, DBNL, and DBN-1 in *C. elegans*) is an adaptor protein that mediates between the actin cytoskeleton and proteins of different intracellular pathways ^20, 21, 22, 23, 24^. Drebrin-like protein is, for example, an important component of the endocytic machinery. Being selectively recruited at the late stage of the formation of clathrin-coated pits, it directly interacts with dynamin, mediating vesicle scission ^25, 26, 27^. Elimination of the mAbp1 in mice results in moderate reduction of both receptor-mediated and synaptic endocytosis and in a severe impairment of synaptic vesicle recycling. This presynaptic defect is associated with restricted physical capabilities and disturbed neuromotoric behavior of the animals. In particular, they display ruffled up fur, muscle trembling, convulsions, partial paralysis of the hind limbs, and strongly reduced fine motor coordination ^28^. The molecular pathways underlying these defects are not clear.

*C. elegans* drebrin-like protein DBN-1 was first described as a sarcomere component of the body-wall muscle cells regulating actin filament dynamics during muscle contraction ^29^. To further analyze the function of DBN-1, we here generated a new *dbn-1* allele (*vit7)* which harbors a premature stop codon within the ADF-H domain of *dbn-1*. Surprisingly, worms bearing the *vit7* mutation acquired larger than usual body curvature variations resulting in unusually strong bending during locomotion. We succeeded in tracking down the molecular mechanisms of the hyper-bending phenotype of the *dbn-1(vit7)* mutant, and identified *C. elegans* drebrin-like protein DBN-1 as an essential regulator of NCA cation leak channel activity.

## Results

### The *vit7* is a loss-of-function mutation

To generate the new *dbn-1* allele, designated *vit7*, we applied Crispr/Cas9 technology of genome editing. The target was chosen within the third exon to be as close as possible to the start codon within the coding sequence and to contain a GG motif adjacent to the PAM sequence to enhance successful genome editing ^30^. The edited allele was designated *vit7*. Sequencing of *dbn-1(vit7)* genomic DNA confirmed precise targeting and showed a 16 base deletion within the third exon of the K08E3.4 gene (Fig. 1 A and B). This deletion resulted in an open-reading-frame shift, leading to the appearance of a premature stop codon. If the mutant protein were stably expressed *in vivo*, it would consist of the 134 N-terminal amino acids of DBN-1, containing the largest part of the ADF-H domain (Fig. 1 C). To analyze protein expression in *vit7*, we generated guinea pig polyclonal antibodies specific against the ADF-H domain of DBN-1 and performed western blot analysis of mutant worm lysate. While these antibodies recognized the ADF-H domain of DBN-1 expressed from an extrachromosomal array, the corresponding band of truncated DBN-1 in the *vit7* mutant was extremely weak or not detectable (Fig. 1 D). This result suggests that truncated DBN-1 was expressed in *vit7 in vivo*, but appeared to be not stable and prone to degradation. Based on this result, we classify *vit7* as a severe loss-of-function allele.

**Figure 1.**
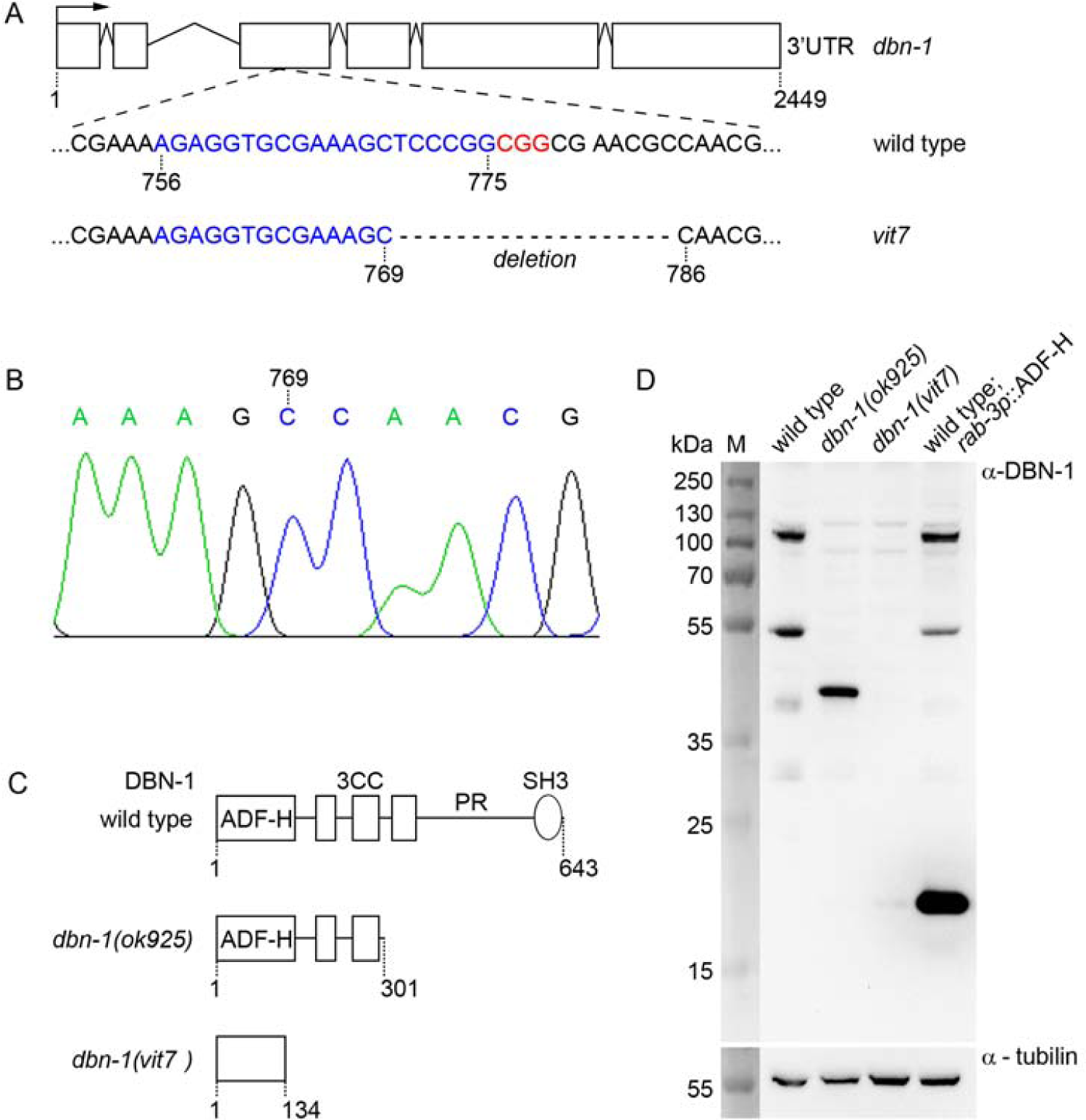
Schematic representation of the *dbn-1(vit7)* allele and protein expression in the *dbn-1(vit7)* mutant. (A) Schematic representation of the *dbn-1* gene. Blue indicates the sgRNA target sequence, and red the PAM motif. The *vit7* allele has a 16-nucleotide deletion within the third exon. (B) Sanger sequencing of genomic DNA of the *vit7* mutant confirms the CRISPR-induced mutation. (C) Schematic representation of DBN-1 expressed in *dbn-1(vit7)* compared to wild type and *dbn-1(ok925)*. (D) SDS western blot analysis of DBN-1 expression in *dbn-1(vit7)* compared to wild type, *dbn-1(ok925)*, and Ex[rab-3p∷ADF-H] worms, using antibodies raised against the ADF-H domain of DBN-1. Molecular mass markers (M) in kD are indicated on the left.

### The *vit7* mutant displays a hyper-bending locomotion phenotype

When crawling on laboratory plate, wild-type *C. elegans* displays a characteristic sinusoidal movement pattern (Fig. 2, A). The wavelength and amplitudes of body bends are highly conserved quantities among wild-type *C. elegans* individuals, across mutants, and across different species ^31, 32^. The *dbn-1(vit7)* mutant worms could be easily distinguished by their strong body curvature and hyper-bending locomotion (Supplementary Movie 1). We recorded and analyzed forward movements of individual worms in the L4 stage by video microscopy and image processing (Fig. 2). When placed on a fresh nematode growth media (NGM) plate with a thin lawn of *E. coli* OP50, *vit7* mutants moved displaying 17.4 ± 3.6 body bends in 20 seconds, significantly less than wild type worms which showed 22.2 ± 4.4 body bends in the same time interval (P < 0,001) (Fig. 2 B). The *vit7* mutants advanced forward on their tracks on average by about one third of their body length per second (with a maximum velocity of half a body length per second) similar to wild type (Fig. 2 C, 4 C, 5 C, 6 C). Mutant animals did thus neither appear to be significantly sluggish nor hyperactive, and their decreased body bend rate resulted rather from hyper-bending. To quantitatively characterize the bending motions of the mutants, we analyzed their body curvature parameters during movement (Fig. 2 D). The mean squared curvature of the body axis of *vit7* mutants (1.78 ± 0.4 × 10^−4^ 1/μm^2^) was two times larger than that of the wild type (0.9 ± 0.2 × 10^-4^ 1/μm^2^) (Fig. 2 E). This result leads us to conclude that DBN-1 is involved in the control of body curvature of *C. elegans*, limiting their degree of bending during locomotion.

**Figure 2.**
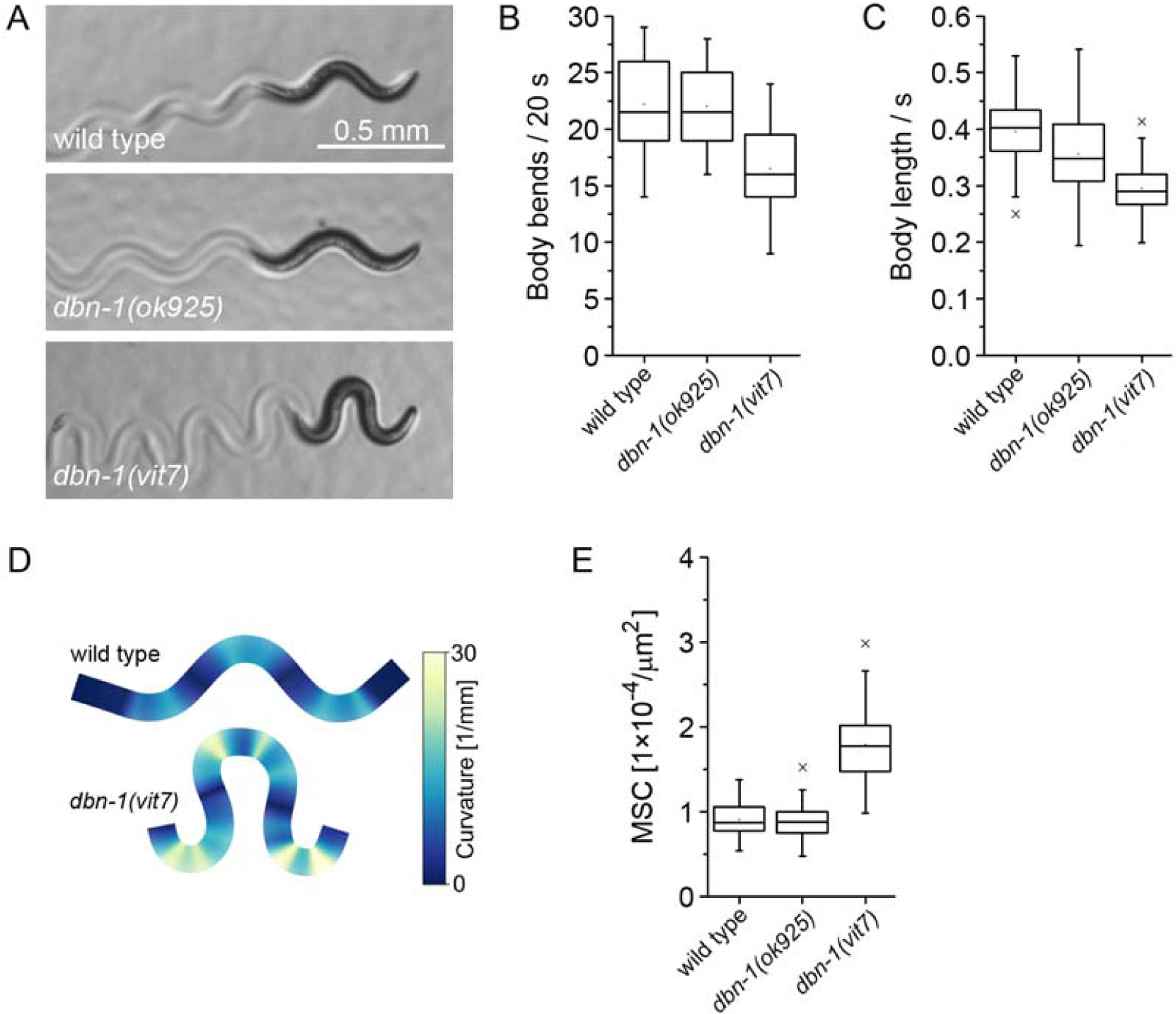
The *dbn-1(vit7*) mutant displays a hyper-bending locomotion phenotype. (A) Images of L4 wild type, *dbn-1(ok925)* and *dbn-1(vit7)* worms. The *dbn-1(vit7)* mutant shows increased body curvature and hyper-bended traces. (B) Quantification of movement velocity as number of body bends in 20 seconds (N = 40). (C) Quantification of movement velocity as body length displacement in one second (N = 50). (D) Graphic representation of the body curvature of *dbn-1(vit7)* mutant in comparison to wild type. (E) Quantification of mean squared curvature (N = 50).

**Figure 3.**
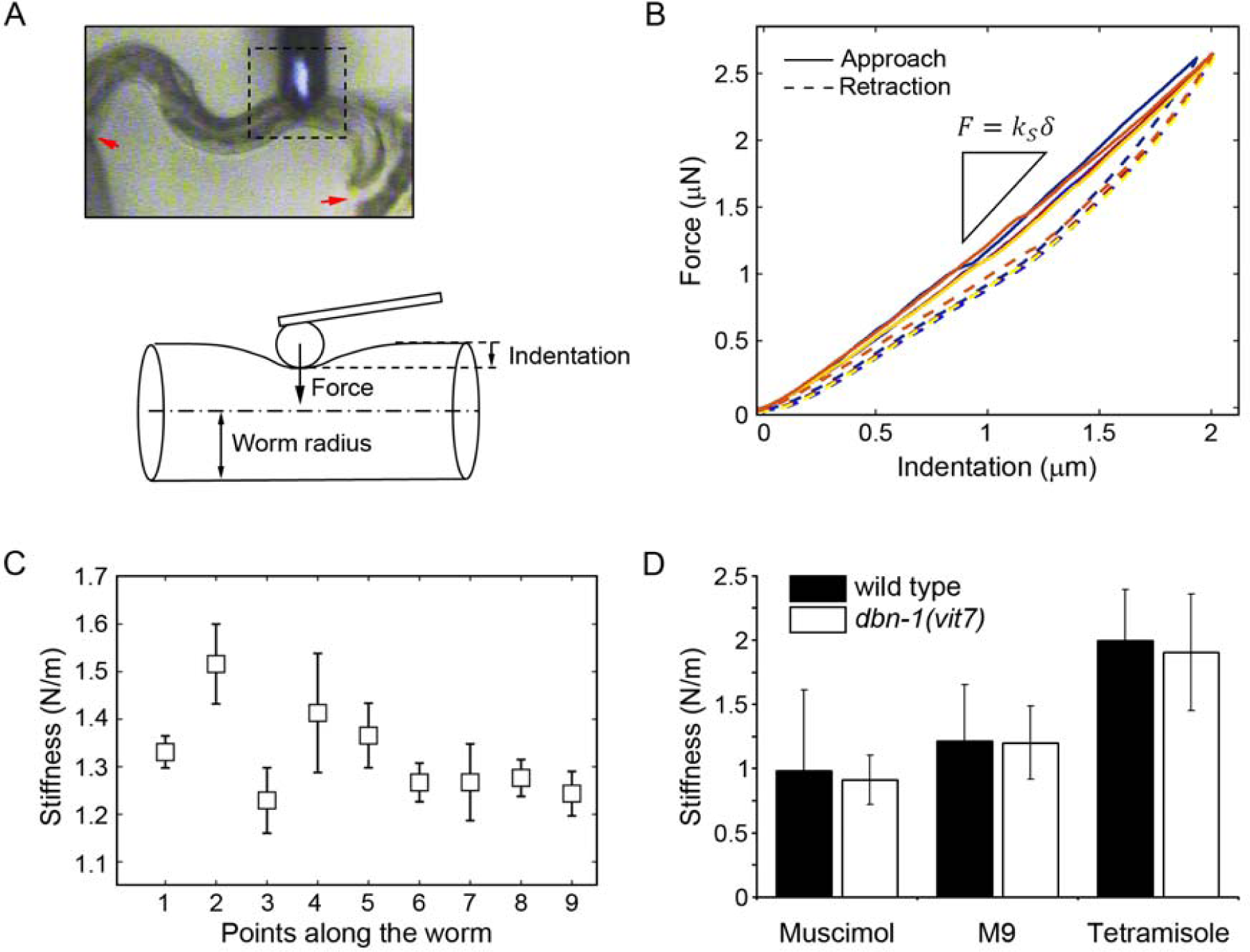
The *dbn-1(vit7)* mutant maintains normal muscle tone on average. (A) Upper panel: Top view of a worm indented with an AFM cantilever (dotted line). The worm was immobilized by gluing the head and tail to the surface of the agar plate (red arrows). Lower panel: schematics of the experiment in a lateral view. The worm was indented with a bead of 7 µm diameter attached to an AFM cantilever. (B) Force-indentation curves in a wild type (untreated) worm. Mechanical response of the nematode remained unchanged over ten subsequent measurements on the same spot (five curves are shown for clarity). Speed of tip approach and retraction was 10 µm/s. The stiffness of the worm (*ks*) was obtained through a linear fit to the approach data in the range of 0.5 to 1.5 µm indentation depth (*δ*). (C) Average stiffness of different spots along a wild type nematode. Each point represents the average of ten measurements, error bars are given by two times the standard deviation. (D) The wild type and *vit7* worms have similar stiffness at physiological conditions in M9 buffer (N = 17 and 21, P = 0.9). Upon treatment with muscimol, the stiffness of wild type and *vit7* worms decreased by 18% (N = 19, P = 0.1) and 24% (N = 24, P = 0.0002) respectively. Treatment with tetramisole resulted in an increase of stiffness of 66% for wild type (N = 27, P = 0.0001) and 59 % for *vit7* (N = 20, P = 0.0001). Values are means, error bars are standard deviations.

**Figure 4.**
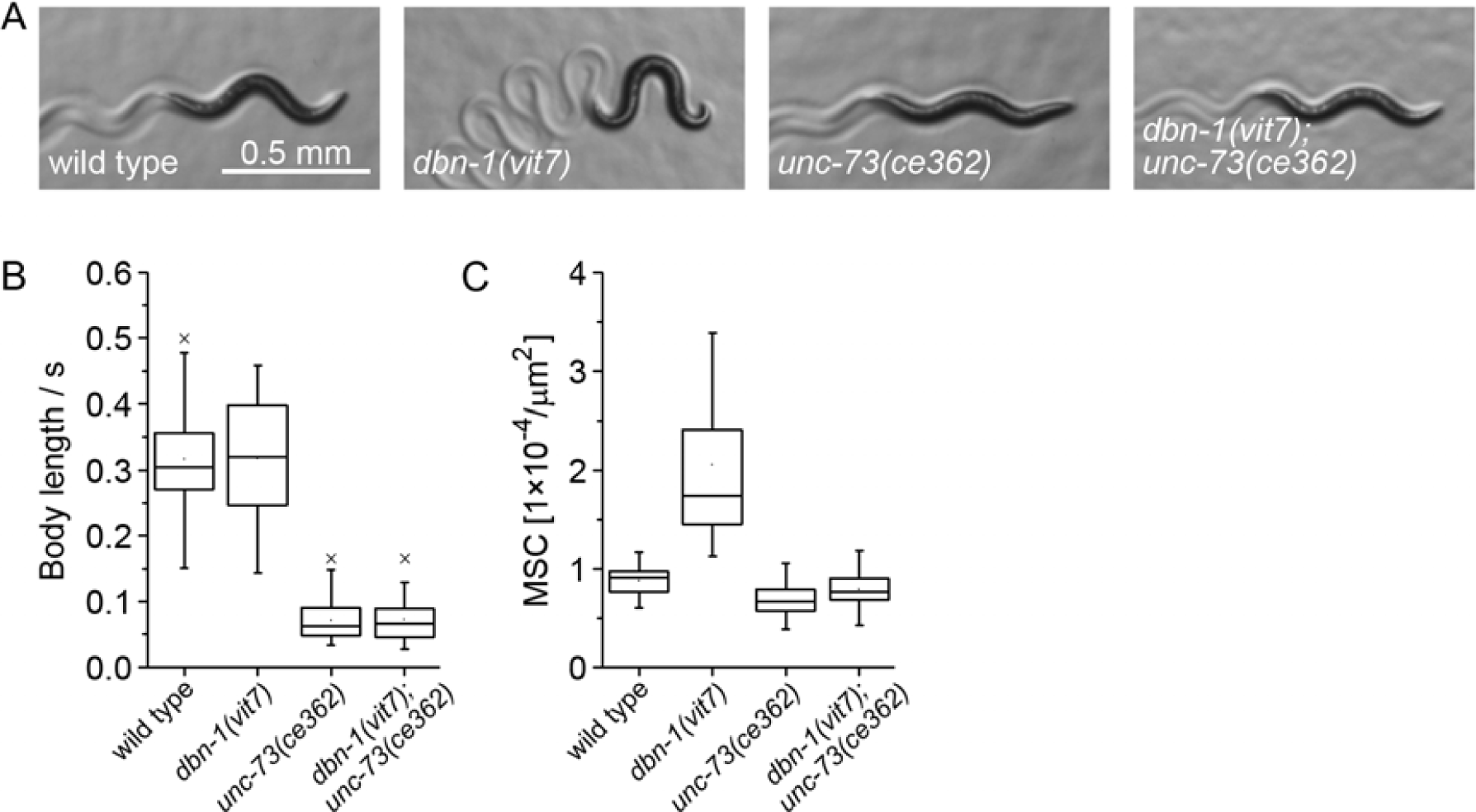
Mutation of Rho-GEF *unc-73(ce362)* suppresses the velocity and hyper-bending phenotype of the *dbn-1(vit7) mutant*. (A) Images of L4 stage wild type worm, *dbn-1(vit7), unc-73(ce362)* and *dbn-1(vit7); unc-73(ce362)* mutants. (B) Quantification of movement velocity. The *unc-73(ce362)* mutation decreases movement velocity of *dbn-1(vit7)* mutant worms (N = 50, P < 0.0001). (C) Quantification of mean squared curvature. The *unc-73(ce362)* mutation suppresses the hyper-bending phenotype of the *dbn-1(vit7)* mutant worms (N = 50, P < 0.0001).

**Figure 5.**
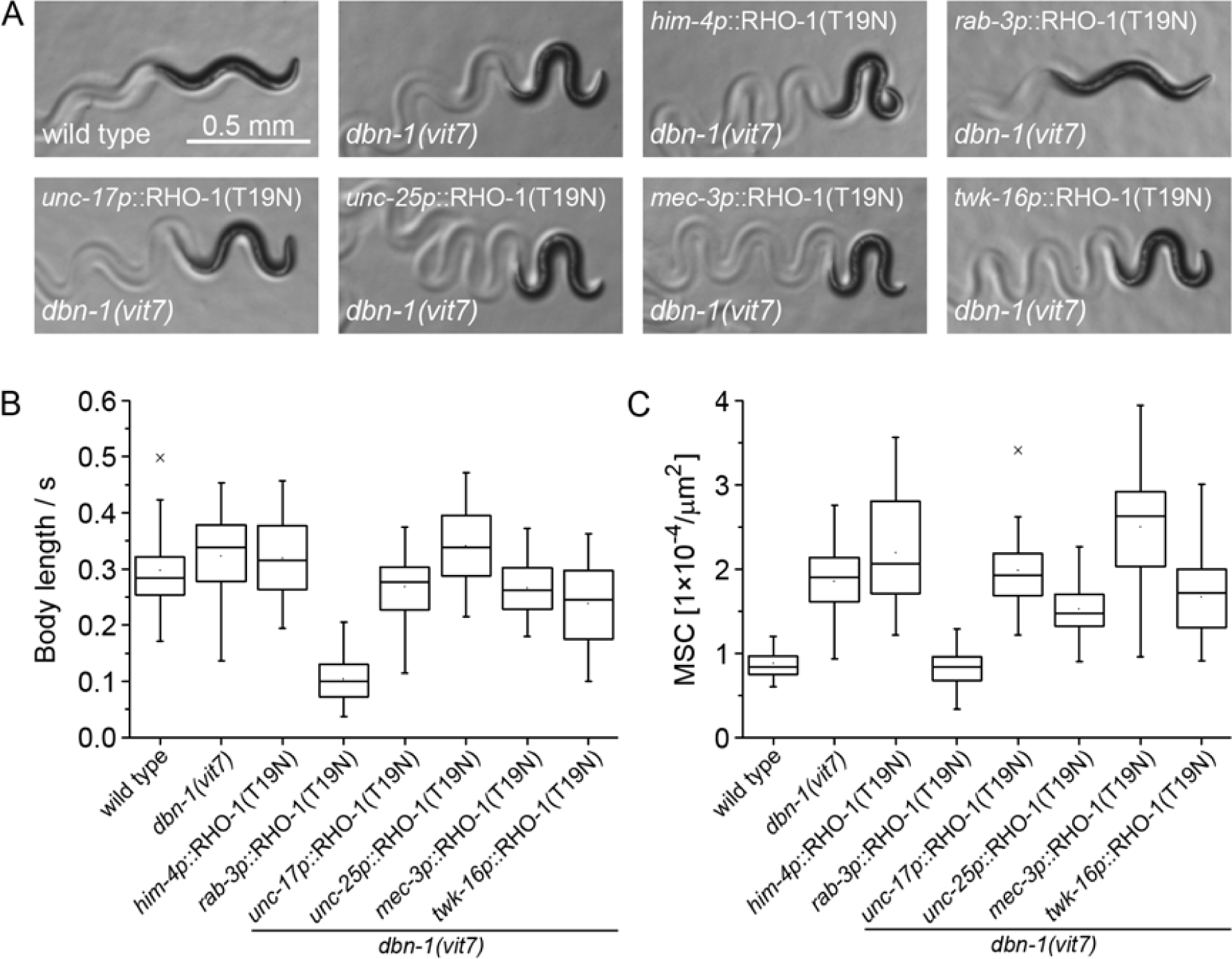
Pan-neuronal expression of dominant negative RHO-1 decreases *dbn-1(vit7)* velocity and suppresses its hyper-bending phenotype. (A) Images of L4 stage wild type worm, *dbn-1(vit7)* and *dbn-1(vit7)* expressing dominant negative RHO-1(T19N) in body-wall muscles (*him-4p*), all neurons (*rab-3p*), acetylcholine neurons (*unc-17p*), GABA neurons (*unc-25p*), mechanosensory neurons (*mec-3p*) and interneurons including DVA neuron (*twk-16p*) from extrachromosomal arrays. (B) Quantification of movement velocity. Expression of *rab-3p*∷RHO-1(T19N) significantly decreases movement velocity of *dbn-1(vit7)* mutant (N = 50, P < 0. 0001). (C) Quantification of mean squared curvature. Expression of *rab-3p*∷RHO-1(T19N) significantly suppresses the hyper-bending phenotype of the *dbn-1(vit7)* mutant (N = 50, P < 0.0001).

**Figure 6.**
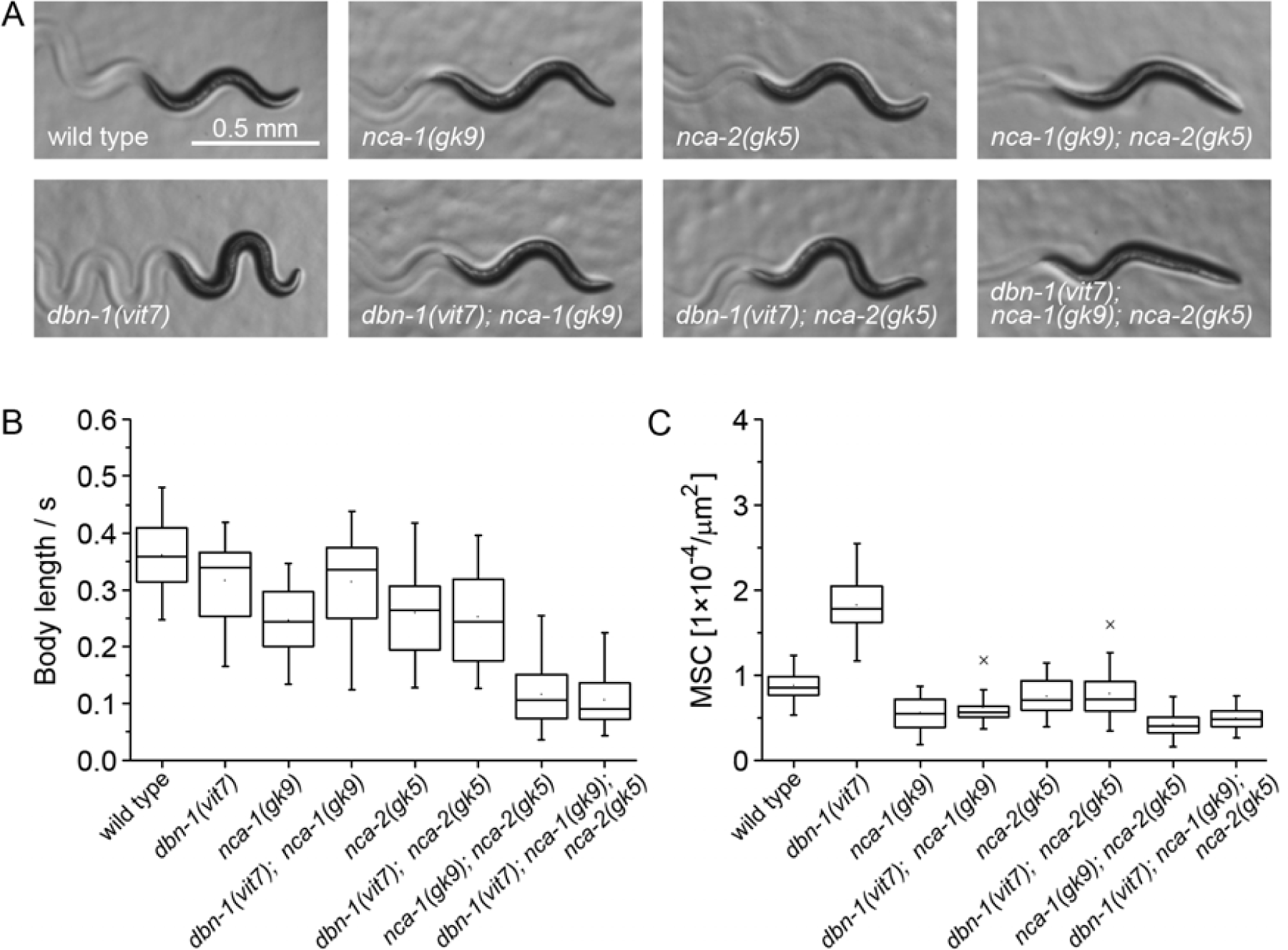
Mutations in NCA-1 and NCA-2 channels suppress the hyper-bending phenotype of the *dbn-1(vit7)* mutant. (A) Images of L4 stage wild type, *dbn-1(vit7), nca-1(gk9), dbn-1(vit7); nca-1(gk9), nca-2(gk5), dbn-1(vit7); nca-2(gk5), nca-1(gk9); nca-2(gk5), dbn-1(vit7); nca-1(gk9); nca-2(gk5)* worms. (B) Quantification of movement velocity. (C) Quantification of mean squared curvature. Single mutations in *nca-1* and *nca-2* suppress the hyper-bending phenotype of the *dbn-1(vit7)* mutant (N = 50, P < 0. 0001). Mutations in both *nca* channels convert the *dbn-1(vit7)* mutant to the “fainter” phenotype, indistinguishable from *nca-1(gk9); nca-2(gk5)* (N = 50, P > 0.5).

**Supplementary Movie 1**. Movement of *C. elegans dbn-1(vit7)* mutant on an NGM plate covered with a thin layer of *E. coli* OP 50 recorded for 20 s, with a rate of 13 frames per second.

### The *vit7* mutant maintains normal average muscle tone

We hypothesized that the hyper-bending phenotype of the *dbn-1(vit7)* mutant might be associated with a changed muscle tone. To test this hypothesis, we used atomic force microscopy to quantify the elastic resistance of the shell-like muscular envelope of the worm body to lateral indentation with a blunt tip. To perform this experiment, we immobilized single worms onto an agar pad by gluing their heads and tails to the surface and then gently indented their bodies with a micron-sized bead attached to an AFM cantilever (Fig. 3 A). The average stiffness of the worm’s muscle shell was calculated from measurements at several different points along the worm’s body. Using a relatively small bead, compared to the dimensions of the worm (tip radius 7 µm << radius of the worm) ensures that the measured resistance is dominated by the outer shell (composed of the cuticle, the hypodermis and the body-wall muscles) and not by an increase of the inner pressure of the worm ^33^. The worms typically lie on their side which means that a probe applied from above test both dorsal and ventral muscles. Furthermore, the time resolution of the experiment was about 5 s, meaning that fast fluctuations of muscle tone were averaged over. The method is thus useful for determining time- and spatially averaged muscle tone. To test the method, we used drug interference to either relax or persistently tense the body-wall muscles. To relax, worms were treated with the GABA receptor agonist muscimol (20 mM), persistent muscle contraction was induced by treatment with the acetylcholine receptor agonist tetramisole (5 mM). The amount of hysteresis seen from the difference in approach and retraction curves (Fig. 3 B) was small, indicating that the response to indentation is dominated by the elasticity of the tissues. Reproducibility in a given location on the worm body was within 1-5% (Fig. 3 B). Response varied at the very ends of the worms’ bodies and over the embryos. These locations were therefore excluded. Variability along the body axis and between different worms was within 30% and 40% respectively (Fig. 3 C).

The *vit7* and wild type worms showed similar stiffness at physiological conditions in control buffer (1.2 ± 0.28 N/m, N = 21 for *vit7* vs 1.21± 0.43 N/m, N = 17 for wild type). Upon treatment with muscimol, the stiffness of *vit7* and wild type worms decreased by 24% and 18% respectively (0.91 ± 0.19 N/m, N = 24 for *vit7* vs 0.98 ± 0.36, N = 19 for wild type). Treatment with tetramisole resulted in an increase in stiffness by 59 % for *vit7* and 66% for wild type (1.9 ± 0.45 N/m, N = 20 for *vit7* vs 1.99 ± 0.4 N/m, N = 27 for wild type) (Fig. 3 D). The measured resistance to indentation of both *dbn-1(vit7)* and wild type worms was thus clearly correlated with the level of muscle contraction. However, the average body-wall-muscle rigidity of *vit7* mutants was not significantly different from that of wild type worms. This result indicates that *dbn-1(vit7)* mutant is able to maintain proper muscle tone on average.

### Inhibition of Rho suppresses the *vit7* mutant phenotype

Aiming at tracking down the molecular mechanism of hyper-bending of the *vit7* mutant, we hypothesized that DBN-1 might be a novel regulator of the recently discovered signaling cascade from Gq via RHO-1 to NCA ion channels that regulates neuronal excitability and synaptic release ^17^. The motivation for this hypothesis was that it was found, first, that *C. elegans* mutants with activated Gq pathway display coiled posture and a “loopy” waveform of motion ^34, 17^. Mutations of players downstream of Gq, namely RhoGEF, NCA channels and NCA channel accessory subunits (UNC-79, UNC-80 and NLF-1) have been shown to suppress the effect of activated RHO-1 ^34, 17^. Second, depletion of mammalian drebrin-like protein mAbp1 or ectopic expression of its ADF-H domain increase Rho GTPase signaling in human breast cancer cells ^24^. Therefore, it is reasonable to assume that the *dbn-1(vit7)* mutation in *C. elegans* might over-activate RHO-1, leading to the observed hyper-bending locomotion.

To test this hypothesis, we studied mutants in proteins downstream of Gq. *C. elegans* carrying the *ce362* mutation has a strongly suppressed RhoGEF function of UNC-73 and cannot be activated by Gq ^34^. We constructed double mutants *dbn-1(vit7); unc-73(ce362)* and found that *ce362* decreased body curvature of the double mutant to the level of an *unc-73(ce362)* single mutant. Moreover, the *dbn-1(vit7); unc-73(ce362)* double mutant appeared sluggish, similar to the *unc-73(ce362)* single mutant (Fig. 4). This result shows that DBN-1 is indeed involved in the regulation of the Gq - Rho signaling pathway opposing RHO-1.

### The locomotion defect of the *vit7* mutant has a neuronal origin

DBN-1 is expressed in multiple tissues including neurons and body-wall muscles (Supplementary Data 1). We therefore needed to identify the tissue in which DBN-1 functions to control body bending. As *C. elegans* has a single gene encoding for Rho, we expressed a dominant negative form of RHO-1 either in body-wall muscles or in neurons of the *dbn-1(vit7)* mutant. RHO-1(T19N), expressed in body-wall muscles (driven by the *him-4 promoter*), did not affect the body curvature of the *dbn-1(vit7)* mutant, whereas its pan-neuronal expression (*rab-3 promoter*) resulted in the complete suppression of the hyper-bending phenotype and in an additional suppression of movement velocity of the *vit7* mutant (Fig. 5). Based on these results, we conclude that DBN-1 controls *C. elegans* body bending from neurons and that RHO-1 has an extended function in the control of movement velocity.

To narrow down which neurons are crucial for DBN-1-mediated body bending control, we expressed RHO-1(T19N) in acetylcholine neurons (driven by the *unc-17 promoter*), GABA neurons (*unc-25 promoter*), mechano-sensory neurons (*mec-3 promoter*) and interneurons including the DVA neuron (*twk-16 promoter*) of the *vit7* mutant. Remarkably, inactivation of RHO-1 in none of these classes of neurons alone resulted in the suppression of the hyper-bending phenotype of *vit7* mutant (Fig. 5). The mechanism regulating body curvature is therefore not exclusively located in either acetylcholine, GABA, mechano-sensory or interneurons.

### Mutations in NCA-1 and NCA-2 suppress the *vit7* mutant phenotype

Finally, we tested the hypothesis that DBN-1 is functionally engaged in the regulation of NCA sodium leak channels. NCA-1 and NCA-2 function redundantly: while *nca-1(gk9)* and *nca-2(gk5)* single mutants have wild-type-like sinusoidal wave locomotion, a double *nca-1(gk9); nca-2(gk5)* mutant displays a “fainter” phenotype, characterized by short periods of movement interrupted by periods of inactivity ^35^. Mutations that eliminate NCA channels suppress the locomotion and body posture phenotypes of an activated Rho mutant ^17^. Conversely, gain-of-function *nca* mutants with constitutively open channels show coiled postures and move with strong bending ^8, 36, 17^.

Therefore, if the *vit7* mutation indeed activates RHO-1 signaling within the Gq-NCA pathway, the hyper-bending phenotype of the *vit7* mutant should be suppressed by elimination of NCA channels. We generated two double mutants *dbn-1(vit7); nca-1(gk9)* and *dbn-1(vit7); nca-2(gk5)* as well as the triple mutant *dbn-1(vit7); nca-1(gk9); nca-2(gk5)* and analyzed their locomotion phenotypes. Additional mutations in *nca-1(gk9)* and in *nca-2(gk5)* both suppressed the hyper-bending phenotype of the *dbn-1(vit7)* mutant to the level of single *nca-1(gk9)* and *nca-2(gk5)* mutants. Elimination of both NCA channels in the *dbn-1* mutant converted it to the “fainter” phenotype. The body curvature of the *dbn-1(vit7); nca-1(gk9); nca-2(gk5)* triple mutant was indistinguishable from an *nca-1(gk9); nca-2(gk5)* double mutant (Fig. 6). These results clearly demonstrate that the hyper-bending phenotype of *dbn-1(vit7)* mutant is caused by the enhanced activity of both NCA-1 and NCA-2 channels.

Taken together, our data shows that DBN-1 is involved in neuronal control of *C. elegans* locomotion, limiting the degree of body bending. At the molecular level, DBN-1 opposes RHO-1 and suppress the activity of both NCA-1 and NCA-2 cation leak channels.

## Discussion

In this study, we generated a new *dbn-1(vit7)* mutant allele, which displays a hyper-bending locomotion phenotype, but is able to maintain wild type muscle tone on average. We tracked down the molecular mechanism of this phenotype and identified DBN-1 as an essential regulator of NCA channel activity in neurons (Fig. 7).

**Figure 7.**
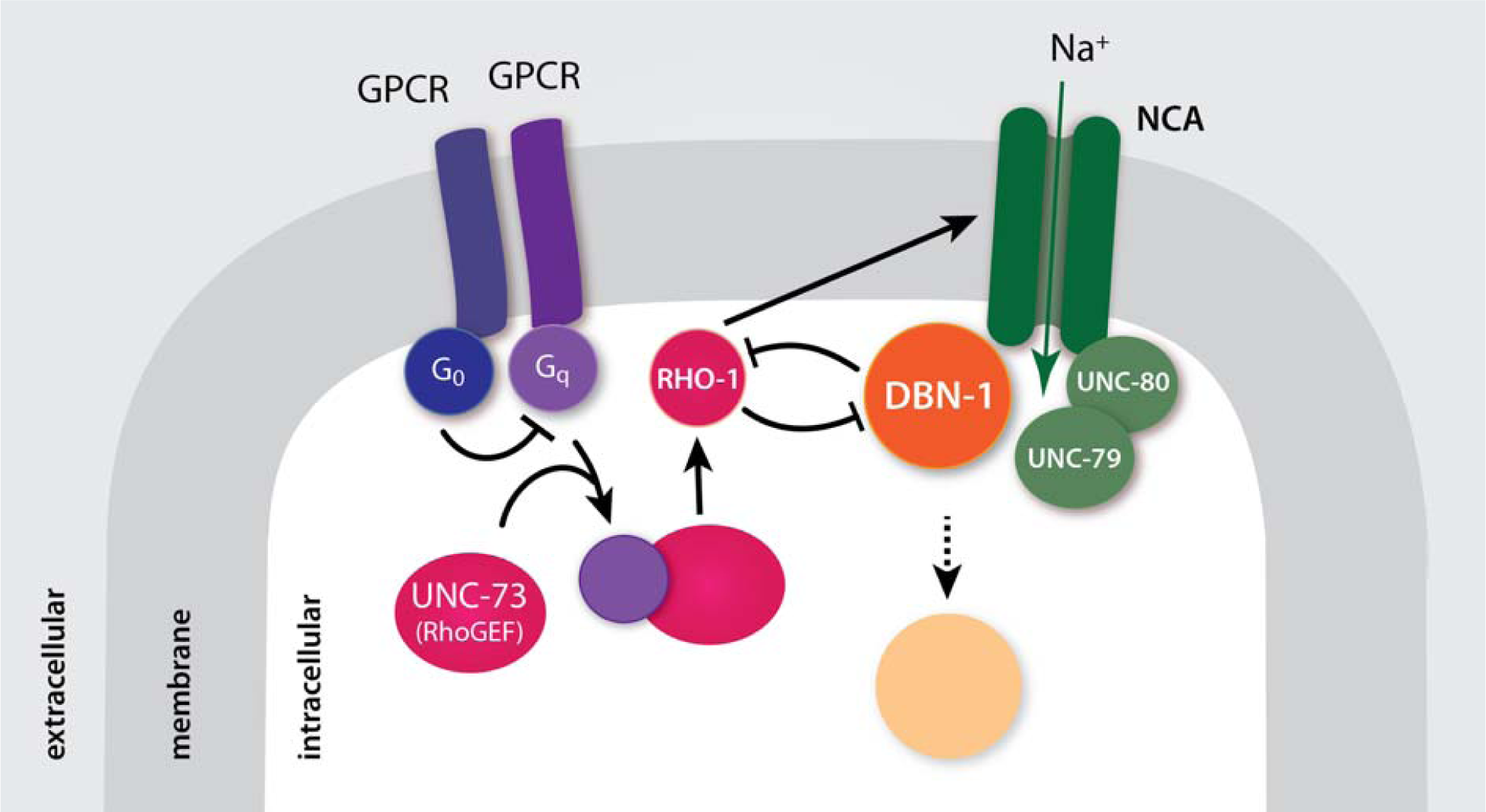
Model of DBN-1 function within the non-canonical Gq pathway. Gq directly binds and activates UNC-73/RhoGEF, which subsequently activates RHO-1 and NCA channel function ^17^. DBN-1 functions antagonistically to RHO-1. DBN-1 targets NCA channels for endocytosis.

*C. elegans* DBN-1 is a multi-domain protein that contains an N-terminal ADF-H domain, three coiled-coils (3xCC), a proline-rich sequence and a C-terminal SH3 domain, of which the 3xCC domain of DBN-1 was shown to bind F-actin *in vitro* ^29^. The *ok925* mutation that truncates DBN-1 after two coiled-coils leads to a mild disorganization of actin filaments during body-wall muscle contraction ^29^ and affects vesicle scission during endocytosis in the intestine ^27^. However, the remaining N-terminal part of DBN-1 in the *ok925* mutant is sufficient to support its wild-type-like locomotion (Fig. 2). In contrast, the truncation of DBN-1 after the largest part of ADF-H domain in the *vit7* mutant we present here dramatically changed the *C. elegans* movement phenotype. This phenotype could be caused either by the absence of the DBN-1 function or by the newly acquired function of the truncated DBN-1 that is expressed at a low level. It would be interesting to know whether coiled-coil domains provide functional redundancy for DBN-1 in actin binding. Whether or not this is the case, it is evident that the 3xCC domain of DBN-1 plays an essential role in the molecular pathway controlling the body bending mechanism.

Recently, Boateng and colleagues discovered that the mammalian drebrin-like protein mAbp1 counteracts Rho GTP-ase in breast cancer cells ^24^. Our data clearly indicate that an analogous mechanism is at work in *C. elegans*, because both the RhoGEF suppression and the pan-neuronal expression of the dominant negative RHO-1 decreased the body curvature and the movement velocity of the *dbn-1(vit7)* mutant. Interestingly, the expression of RHO-1(T19N) in different classes of neurons (acetylcholine, GABA, mechano-sensory and interneurons including the DVA neuron), as well in the body-wall muscles, did not affect the locomotion behavior of *dbn-1(vit7)*. Thus, DBN-1 acts in all neurons to regulate body curvature. This is consistent with the finding that the inhibition of Rho in all neurons suppresses the “loopy” waveform of the activated Gq mutant ^17^.

We here show that DBN-1 suppresses the activity of NCA sodium leak channels. NCA-1 and NCA-2 are important players that control the neuronal excitability and synaptic transmission at neuromuscular junctions ^8, 9^. The elimination of NCA channels results in hyperpolarized resting membrane potential in premotor interneurons ^10^. The *nca-1; nca-2* loss-of-function mutants expose a “fainter” phenotype, characterized by short periods of movement interrupted by periods of inactivity ^35^. Conversely, gain-of-function *nca* mutants with constitutively open channels show coiled postures and move with strong bending ^8, 36, 17^. While the multiple signaling cascades controlling the NCA channels activity have been identified, is not known how their spatial and temporal regulation is achieved. We speculate that the NCA conductance is regulated by factors that determine the number of active channels at the cell surface, in a similar way as described for other ion channels ^37, 38, 39^. Based on the well-established roles of drebrin-like protein in endocytosis ^25, 40^, ^28, 26, 27^, it seems likely that interaction of DBN-1 with NCA channels directs their internalization. In support of this hypothesis, mAbp1 knockout mice display a moderate reduction of both receptor-mediated and synaptic endocytosis as well as severe impairment in synaptic vesicle recycling ^28^, whereas elimination NCA channels ameliorates the recycling defect in synaptojanin mutants, increasing the number of synaptic vesicles in both GABA and acetylcholine neurons ^41^. In addition, mAbp1 has been discovered to be a key component of the immunological synapses that regulates surface clustering of B-cell receptors ^42^. In contrast, Rho is known to inhibit clathrin-mediated endocytosis ^43^. Therefore, the activation of NCA channels by RHO-1 ^17^ might be a result of inhibition of their internalization.

In humans, mutations of NALCN channels are the direct causative factors of IHPRF (infantile hypotonia with psychomotor retardation and characteristic facies) and CLIFAHDD (congenital contractures of the limbs and face with hypotonia and developmental delay) syndromes. While the loss-of-function of NALCN leads to variable degree of hypotonia ^44^, the most striking clinical abnormality of NALCN hyperactivity is a markedly delayed relaxation of muscles following activation ^36^. We measured the muscle response of *dbn-1(vit7)* mutants using atomic force microscopy. In spite of the fact that the animals have an obvious defect in muscle regulation, we could not detect significant disturbance in their average muscle tone. The reason for this might be that the *dbn-1(vit7)* mutant has a less severe phenotype than the *C. elegans* bearing a orthologous patient’s *NALCN* mutation, which leads to curly posture, small body size, and reduced locomotion ^36^.

In summary, this study has revealed a surprising role of DBN-1 in the molecular mechanisms that control the degree of body bending of *C. elegans* via suppression of NCA sodium leak channels. We suggest that the level of dynamic activity of NCA channels that is required for sustained locomotion is set by the balance between membrane insertion and recycling. Because the structures of ion channels, receptors and their regulatory machineries are remarkably conserved between *C. elegans* and humans ^45^, a similar mechanism might be at work in the regulation of human NALCN conductance. The *dbn-1(vit7)* mutant therefore appears to be a suitable model to study the molecular mechanisms and pathways underling NALCN channel function.

## Materials and methods

### C. elegans strains

Wild type Bristol N2 and mutant strains: *unc-73(ce362), nca-1(gk9), nca-2(gk5)* and *nca-1(gk9); nca-2(gk5)* were obtained from the *C. elegans* Genetic Center (University of Minnesota). Worms were cultured at 20°C on nematode growth media (NGM) plates seeded with *E. coli* OP50 ^46^.

### Generation of the dbn-1(vit7) mutant

The design of sgRNA was performed using CRISPRdirect software ^47^. To achieve successful genome editing ^30^, a target-specific sequence 5’-GAGGTGCGAAAGCTCCCGG-3’ was chosen, containing a GG motif adjacent to the PAM sequence. The 19 base-sgRNA was cloned into pDD162 using a Q5 mutagenesis kit (New England Biolabs), as described by ^48^. pDD162 (Peft-3∷Cas9 + Empty sgRNA) ^49^ was purchased from Addgene (Plasmid #47549). A repair oligo containing mutated PAM and new bases inserted between the sgRNA sequence and PAM (5’-CATGCCAGATCGGAAATCGACATCGAGCCGGACGCGATTCGAAAAGAGGTGCGAAAGCTCCCGGGCTAGCTCGTCGAACGCCAACGCCATCGTCGAATCCACGTACAGCATGCCGGAG-3’) was synthesized by Sigma-Aldrich. A mixture of Cas9/sgRNA1 for DBN-1 (50 ng/μl), repair oligo (30 ng/μl) and co-injection marker pRF4 (rol-6(su1006)) (120 ng/μl) was injected into the gonads of *C. elegans* ^50^. Edits were recovered from F2 plate containing rollers. To verify genome editing, genomic DNA encoding DBN-1 was amplified by PCR, cloned into the pDonr201 vector (Invitrogen) and sequenced by Seqlab. Although a repair oligo was designed to insert a stop codon, the genome change resulted instead in a 16 base pairs deletion, which caused an open reading frame shift and led to the appearance of a premature stop codon. Only a single *vit7* allele was found. To ensure the absence of mutations in other genes, the original mutant strain was back-crossed 5 times with wild type.

### Molecular biology

*Unc-25, unc-17, mec-3, twk-16* (containing a 491 bp sequence upstream of the start codon, first exon and first intron ^51^) and *him-4* promoter sequences were amplified from *C. elegans* genomic DNA and each cloned to replace a *rab-3* promoter in the rab-3p∷GW (gift from Stefan Eimer, University of Frankfurt). The *rho-1* cDNA was amplified from *C. elegans* cDNA (Invitrogen), sub-cloned into a pDonr201 vector (Invitrogen), mutated using the Q5 mutagenesis kit (New England Biolabs) to obtain a T19N mutation and cloned into rab-3p∷GW, unc-25p∷GW, unc-17p∷GW, mec-3p∷GW, twk-16p∷GW and him-4p∷GW destination vectors. A cDNA encoding amino acids 1-132 of DBN-1 was amplified from *dbn-1* cDNA and cloned into pET28a and rab-3∷GW vectors. All cDNA clones were confirmed by DNA sequencing at Seqlab.

### Transgene generation and crossings

Transgenic strains were generated by microinjection ^50^ of the plasmids dbn-1∷GFP-dbn-1 (100 ng/μl), rab-3∷rho-1(T19N) (3 ng/μl), him-4∷rho-1(T19N) (50 ng/μl), unc-17∷rho-1(T19N) (10 ng/μl), unc-25∷rho-1(T19N) (15 ng/μl), mec-3∷rho-1(T19N) (25 ng/μl), twk-16∷rho-1(T19N) (30 ng/μl) into *dbn-1(vit7)* and rab-3p∷ADF-H (10 ng/μl) into wild type gonads. pRFP-odr-1 (100 ng/μl) was used as co-injection marker. Three to five transgenic lines were isolated for each injected plasmid mix and it was confirmed that each set of lines had the same phenotype. Crossings of *unc-73(ce362), nca-1(gk9), nca-2(gk5)* and *nca-1(gk9); nca-2(gk5)* to *dbn-1(vit7)* were performed using classical genetic approaches. Mutations in the double and triple mutants were confirmed by PCR or sequencing.

### Antibodies against the ADF-H domain of DBN-1

Recombinant His-tagged DBN-1(1-132) was expressed in *E. coli* BL21-Gold cells and purified on Ni-NTA Agarose (Macherey-Nagel). Two guinea pigs were immunized at Bioscience with His-DBN-1(1-132) coupled to keyhole limpet hemocyanin. The final antisera were purified separately using His-DBN-1(1-132) bound to NHS-activated Sepharose High Performance (GE Healthcare Life Science). Both purified antibodies were tested in western blot. They similarly recognized the ADF-H domain of DBN-1 in the lysate of Ex[rab-3p∷ADF-H] worms, DBN-1 in the lysates of wild type and *dbn-1(ok925)* worms, and the significantly reduced level of DBN-1 in the lysate of wild type worms fed with RNAi against DBN-1. The reason for the generation of these antibodies was that the antibodies against full-length DBN-1 described in ^29^ do not recognize the ADF-H domain of DBN-1.

### Protein electrophoresis and western blots

Hundred young adult worms were collected in 1.5 ml tubes in 25 μl of lysis buffer, containing 150 mM NaCl, 50 mM Tris pH7.4, 2mM EDTA and 1% Nonidet P-40. The samples were then subjected to alternating procedures of freezing in liquid nitrogen followed by heating at 24° to break the worms. After this, samples were homogenized by pipetting 100 times up and down using a 10 μl pipette tip. 25 μl of 2x Laemmli buffer was added and the sample was heated to 95°C for 5 min. 15 μl of sample were resolved by a 12% NuPAGE Bis-Tris gel (Invitrogen). Proteins were transferred onto an Immobilon-P polyvinylidene difluoride membrane (Millipore) using a Bio-Rad Criterion Blotter. After the transfer, the membrane was blocked with 10% nonfat milk in PBS containing 0.05% Tween 20 for 30 min and incubated with affinity-purified guinea pig anti-ADF-H domain-DBN-1 antibodies diluted at 1:500 at 4°C overnight, followed by treatment with diluted at 1:10000 peroxidase-labeled goat anti-guinea pig IgG (sc-2438). The reactivity was detected with an Amersham ECL Detection Reagent (GE Healthcare Life Science) using an Intas gel imager (Intas Science Imaging Instruments GmbH). The membrane was treated with a buffer containing 2% SDS, 100 mM β-mercaptoethanol, and 62.5 mM Tris-HCl, pH 6.8, at 50°C for 30 min to remove bound probes, and re-probed with mouse monoclonal anti-α-tubulin antibody diluted at 1:1000 (T6199, Sigma-Aldrich) in combination with peroxidase-labeled goat anti-mouse IgG diluted at 1:10000 (A6154, Sigma-Aldrich).

### Behavioral Assays

Groups of *C. elegans*, each containing 10 worms, were assayed at the same environmental parameters. An individual worm was placed onto new NGM plate with a thin layer of *E. coli* OP50. Immediately after the worm adapted to the new environment and started to move forward, its movement was recorded using a Leica MZFLIII microscope equipped with a SPOT Insight camera. Videos (each containing 20 frames, taken with a rate of 1 frame per second) were analyzed using ImageJ ^52^. The worms’ velocity was measured as the number of body bends in 20 seconds and as body length per second.

### Numerical calculation of mean squared curvature

To calculate mean squared curvature, we followed established procedures ^53^. A midline ^32^ was drawn along the entire length of the worm body and analyzed by a custom-written computer program. For each worm, five consecutive positions in an interval of 1 s were analyzed. To estimate the mean squared curvature of the worms, we interpolated hand-drawn worm shapes using a basis spline of degree 6 and an interpolation factor of *n* = 100. Only the central constant-diameter section of the body of the worms was considered. Thin body tips were not taken into account since the curvature tended to vary and get stronger at the tips.

The curvature *κ* at any point along a (one dimensional) curve in two dimensions is defined as the rate of change in tangent direction *θ* of the contour, as a function of arc length *s*:

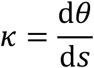

The bending energy of a homogeneous elastic rod with a bending modulus *K* is:

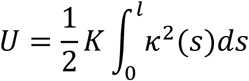

In our case the material properties of the worm are not well known, and it is probably not exactly a homogeneous rod. Nevertheless, we can approximately measure the overall degree of bending *B* by calculating the integral of the squared curvature along the length of the worm’s body:

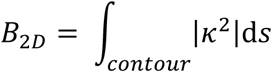

Since worms differed in overall length, we normalized this number with the worm’s contour length *L*, giving us mean squared curvature *MSC* in units of [1/µm^2^]:

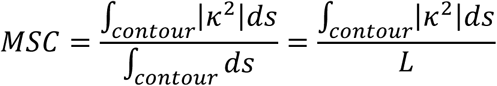

Numerical procedure: The tangent angle *θ* (*s*) at equidistant points along the worm contour (excluding ends) was calculated. The curvature *k*(*s*) was determined as the numerical derivative of *θ* (*s*) with respect to *s*. The worm contour was subdivided into N segments with tangent vector *θ*_i_ and segment length Δ*s*_i_ at each point *i*. The curvature was then calculated as Δ*θ*_i_ *= θ*_i_ − *θ*_i−1_ divided by the segment length Δ*s*_i_:

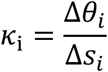

With the total contour length 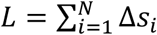, the mean squared curvature was calculated as:

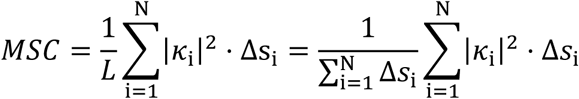

### Body stiffness measurements

Force-indentation curves of living nematodes were captured using an atomic force microscope (MFP-3D, Asylum). Measurements were performed in single wild type and *dbn-1(vit7)* young adult worms either in their physiological state or chemically treated to induce muscle relaxation or muscle contraction. Samples were prepared by placing worms into either M9 buffer or M9 buffer containing 20 mM muscimol (incubation time ∼30-60 min) or 5 mM tetramisole (incubation time ∼ 10-15 min). The worm were then transferred from the respective buffers onto agarose pads (0.5 mm thickness; 10 % agarose in water) and their head and tail were immobilized by placing a small drop of instant glue on top (Loctite, Germany, 401). Worms were then covered with the respective buffers. Indentation tests were performed along the worms’ free midsection (far from glued ends) using a bead of radius 7 µm attached to the tip of a rectangular cantilever (Nano and more TL-NLC-50, nominal stiffness 48N/m). To reduce the effect of geometry-induced variabilities in the measured stiffness, the bead was positioned over the center of the nematode. The maximum applied forces (2-3.5 µN) were set to reach maximum indentation depths in the range of 2-4.5 µm. This is sufficient to indent deeper than the thickness of the cuticle plus the hypodermis (∼0.4 µm). The piezo approach velocity during the indentation was set to 10 µm/s. The stiffness of each worm was obtained by performing ten measurements in quick succession on single points. This procedure was repeated on 5 -10 different points along the body mid axis for each worm. In total, 5 worms were analyzed for each condition. A linear fit to log-log plots of the data not regarding indentations less than 0.5 µm gave an average exponent of *n* = 1.04± 0.06, confirming a linear force-indentation relation. The spring constant of the worm at each measured spot was obtained from a linear fit of the approach part of the force-indentation curves in the range of 0.5-1.5 µm of indentation. The spring constant at each point is the average value of the ten measurements on that point, with the error given by twice the standard deviation from the average.

### Statistical analysis

All statistical tests and computations were carried out using Python (https://www.python.org/) and OriginPro 8.5G (https://www.originlab.com/) software. Values obtained from the worms’ behavioral assays are shown as box-and-whisker plots, where the upper and lower limits of the box correspond to the 75^th^ and 25^th^ percentile values, the central bar indicates the median and the whiskers indicate the minimum and maximum values. Values of the worms’ body stiffness measured by AFM are reported as the mean ± standard deviation. The data were tested for a normal distribution. The group data were compared using a two-tailed Student’s t-test.

## Acknowledgments

We thank Tanja Gall for technical assistance. Some *C. elegance* strains were provided by the CGC, which is funded by NIH Office of Research Infrastructure Programs (P40 OD010440). This research was supported by the European Research Council under the European Union’s Seventh Framework Programme (FP7/2007-2013) / ERC grant agreement n°340528.

## Contributions

E.B. designed the project, performed the molecular biology, biochemical and genetic work, locomotion assay, image processing, data analysis, preparation of figures, and co-wrote the paper. P.W. and R.G performed AFM and analyzed the AFM data. D.R.K. drew the scheme on Fig.7 and provided experimental equipment and advice. D.H wrote a computer program for mathematical analysis of worm’s curvatures. C.F.S. supervised the project and co-wrote the paper.

## Competing financial interests

The authors declare no competing financial interests.

## References

1. Gray, J. & Lissmann, H.W. The Locomotion of Nematodes. The Journal of experimental biology 41, 135–154 (1964).

2. Croll, N.A. Sensory basis of activation in nematodes. Experimental parasitology 27, 350–356 (1970).

3. Von Stetina, S.E., Treinin, M. & Miller, D.M., 3rd The motor circuit. International review of neurobiology 69, 125–167 (2006).

4. Schuske, K., Beg, A.A. & Jorgensen, E.M. The GABA nervous system in C. elegans. Trends in neurosciences 27, 407–414 (2004).

5. Gao, S. & Zhen, M. Action potentials drive body wall muscle contractions in Caenorhabditis elegans. Proceedings of the National Academy of Sciences of the United States of America 108, 2557–2562 (2011).

6. Burr, A.H. & Gans, C. Mechanical significance of obliquely striated architecture in nematode muscle. The Biological bulletin 194, 1–6 (1998).

7. Lu, B. et al. The neuronal channel NALCN contributes resting sodium permeability and is required for normal respiratory rhythm. Cell 129, 371–383 (2007).

8. Yeh, E. et al. A putative cation channel, NCA-1, and a novel protein, UNC-80, transmit neuronal activity in C. elegans. PLoS biology 6, e55 (2008).

9. Gao, S. et al. The NCA sodium leak channel is required for persistent motor circuit activity that sustains locomotion. Nature communications 6, 6323 (2015).

10. Xie, L. et al. NLF-1 delivers a sodium leak channel to regulate neuronal excitability and modulate rhythmic locomotion. Neuron 77, 1069–1082 (2013).

11. Lu, B. et al. Peptide neurotransmitters activate a cation channel complex of NALCN and UNC-80. Nature 457, 741–744 (2009).

12. Swayne, L.A. et al. The NALCN ion channel is activated by M3 muscarinic receptors in a pancreatic beta-cell line. EMBO reports 10, 873–880 (2009).

13. Lu, B. et al. Extracellular calcium controls background current and neuronal excitability via an UNC79-UNC80-NALCN cation channel complex. Neuron 68, 488–499 (2010).

14. Philippart, F. & Khaliq, Z.M. Gi/o protein-coupled receptors in dopamine neurons inhibit the sodium leak channel NALCN. eLife 7 (2018).

15. Bouhours, M. et al. A co-operative regulation of neuronal excitability by UNC-7 innexin and NCA/NALCN leak channel. Molecular brain 4, 16 (2011).

16. Kasap, M., Bonnett, K., Aamodt, E.J. & Dwyer, D.S. Akinesia and freezing caused by Na(+) leak-current channel (NALCN) deficiency corrected by pharmacological inhibition of K(+) channels and gap junctions. The Journal of comparative neurology 525, 1109–1121 (2017).

17. Topalidou, I. et al. The NCA-1 and NCA-2 Ion Channels Function Downstream of Gq and Rho To Regulate Locomotion in Caenorhabditis elegans. Genetics 206, 265–282 (2017).

18. Topalidou, I., Cooper, K., Pereira, L. & Ailion, M. Dopamine negatively modulates the NCA ion channels in C. elegans. PLoS genetics 13, e1007032 (2017).

19. Hoyt, J.M. et al. The SEK-1 p38 MAP Kinase Pathway Modulates Gq Signaling in Caenorhabditis elegans. G3 7, 2979–2989 (2017).

20. Kessels, M.M., Engqvist-Goldstein, A.E. & Drubin, D.G. Association of mouse actin-binding protein 1 (mAbp1/SH3P7), an Src kinase target, with dynamic regions of the cortical actin cytoskeleton in response to Rac1 activation. Molecular biology of the cell 11, 393–412 (2000).

21. Schafer, D.A. et al. Dynamin2 and cortactin regulate actin assembly and filament organization. Current biology : CB 12, 1852–1857 (2002).

22. Fenster, S.D. et al. Interactions between Piccolo and the actin/dynamin-binding protein Abp1 link vesicle endocytosis to presynaptic active zones. The Journal of biological chemistry 278, 20268–20277 (2003).

23. Hou, P. et al. Fgd1, the Cdc42 GEF responsible for Faciogenital Dysplasia, directly interacts with cortactin and mAbp1 to modulate cell shape. Human molecular genetics 12, 1981–1993 (2003).

24. Boateng, L.R., Bennin, D., De Oliveira, S. & Huttenlocher, A. Mammalian Actin-binding Protein-1/Hip-55 Interacts with FHL2 and Negatively Regulates Cell Invasion. The Journal of biological chemistry 291, 13987–13998 (2016).

25. Kessels, M.M., Engqvist-Goldstein, A.E., Drubin, D.G. & Qualmann, B. Mammalian Abp1, a signal-responsive F-actin-binding protein, links the actin cytoskeleton to endocytosis via the GTPase dynamin. The Journal of cell biology 153, 351–366 (2001).

26. He, K. et al. Mammalian actin-binding protein 1/HIP-55 is essential for the scission of clathrin-coated pits by regulating dynamin-actin interaction. FASEB journal : official publication of the Federation of American Societies for Experimental Biology 29, 2495–2503 (2015).

27. Shi, X. et al. WIP-1 and DBN-1 promote scission of endocytic vesicles by bridging actin and Dynamin-1 in the C. elegans intestine. Journal of cell science 132 (2019).

28. Connert, S. et al. SH3P7/mAbp1 deficiency leads to tissue and behavioral abnormalities and impaired vesicle transport. The EMBO journal 25, 1611–1622 (2006).

29. Butkevich, E. et al. Drebrin-like protein DBN-1 is a sarcomere component that stabilizes actin filaments during muscle contraction. Nature communications 6, 7523 (2015).

30. Farboud, B. & Meyer, B.J. Dramatic enhancement of genome editing by CRISPR/Cas9 through improved guide RNA design. Genetics 199, 959–971 (2015).

31. Karbowski, J. et al. Conservation rules, their breakdown, and optimality in Caenorhabditis sinusoidal locomotion. Journal of theoretical biology 242, 652–669 (2006).

32. Nahabedian, J.F., Qadota, H., Stirman, J.N., Lu, H. & Benian, G.M. Bending amplitude - a new quantitative assay of C. elegans locomotion: identification of phenotypes for mutants in genes encoding muscle focal adhesion components. Methods 56, 95–102 (2012).

33. Petzold, B.C. et al. Caenorhabditis elegans body mechanics are regulated by body wall muscle tone. Biophysical journal 100, 1977–1985 (2011).

34. Williams, S.L. et al. Trio’s Rho-specific GEF domain is the missing Galpha q effector in C. elegans. Genes & development 21, 2731–2746 (2007).

35. Humphrey, J.A. et al. A putative cation channel and its novel regulator: cross-species conservation of effects on general anesthesia. Current biology : CB 17, 624–629 (2007).

36. Bend, E.G. et al. NALCN channelopathies: Distinguishing gain-of-function and loss-of-function mutations. Neurology 87, 1131–1139 (2016).

37. Shimkets, R.A., Lifton, R.P. & Canessa, C.M. The activity of the epithelial sodium channel is regulated by clathrin-mediated endocytosis. The Journal of biological chemistry 272, 25537–25541 (1997).

38. Gao, Y., Bertuccio, C.A., Balut, C.M., Watkins, S.C. & Devor, D.C. Dynamin- and Rab5-dependent endocytosis of a Ca2+ -activated K+ channel, KCa2.3. PloS one 7, e44150 (2012).

39. Conrad, R. et al. Rapid Turnover of the Cardiac L-Type CaV1.2 Channel by Endocytic Recycling Regulates Its Cell Surface Availability. iScience 7, 1–15 (2018).

40. Le Bras, S. et al. Recruitment of the actin-binding protein HIP-55 to the immunological synapse regulates T cell receptor signaling and endocytosis. The Journal of biological chemistry 279, 15550–15560 (2004).

41. Jospin, M. et al. UNC-80 and the NCA ion channels contribute to endocytosis defects in synaptojanin mutants. Current biology : CB 17, 1595–1600 (2007).

42. Seeley-Fallen, M.K. et al. Actin-binding protein 1 links B-cell antigen receptors to negative signaling pathways. Proceedings of the National Academy of Sciences of the United States of America 111, 9881–9886 (2014).

43. Lamaze, C., Chuang, T.H., Terlecky, L.J., Bokoch, G.M. & Schmid, S.L. Regulation of receptor-mediated endocytosis by Rho and Rac. Nature 382, 177–179 (1996).

44. Al-Sayed, M.D. et al. Mutations in NALCN cause an autosomal-recessive syndrome with severe hypotonia, speech impairment, and cognitive delay. American journal of human genetics 93, 721–726 (2013).

45. Lai, C.H., Chou, C.Y., Ch’ang, L.Y., Liu, C.S. & Lin, W. Identification of novel human genes evolutionarily conserved in Caenorhabditis elegans by comparative proteomics. Genome research 10, 703–713 (2000).

46. Brenner, S. The genetics of Caenorhabditis elegans. Genetics 77, 71–94 (1974).

47. Naito, Y., Hino, K., Bono, H. & Ui-Tei, K. CRISPRdirect: software for designing CRISPR/Cas guide RNA with reduced off-target sites. Bioinformatics 31, 1120–1123 (2015).

48. Paix, A. et al. Scalable and versatile genome editing using linear DNAs with microhomology to Cas9 Sites in Caenorhabditis elegans. Genetics 198, 1347–1356 (2014).

49. Dickinson, D.J., Ward, J.D., Reiner, D.J. & Goldstein, B. Engineering the Caenorhabditis elegans genome using Cas9-triggered homologous recombination. Nature methods 10, 1028–1034 (2013).

50. Mello, C. & Fire, A. DNA transformation. Methods in cell biology 48, 451–482 (1995).

51. Salkoff, L. et al. Evolution tunes the excitability of individual neurons. Neuroscience 103, 853–859 (2001).

52. Schneider, C.A., Rasband, W.S. & Eliceiri, K.W. NIH Image to ImageJ: 25 years of image analysis. Nature methods 9, 671–675 (2012).

53. Young, I.T., Walker, J.E. & Bowie, J.E. An analysis technique for biological shape. I. Information and control 25, 357–370 (1974).

