## Supplemental Figure 1 for "Drebrin-like protein regulates body bending of *C. elegans* via suppression of NCA cation leak channels"

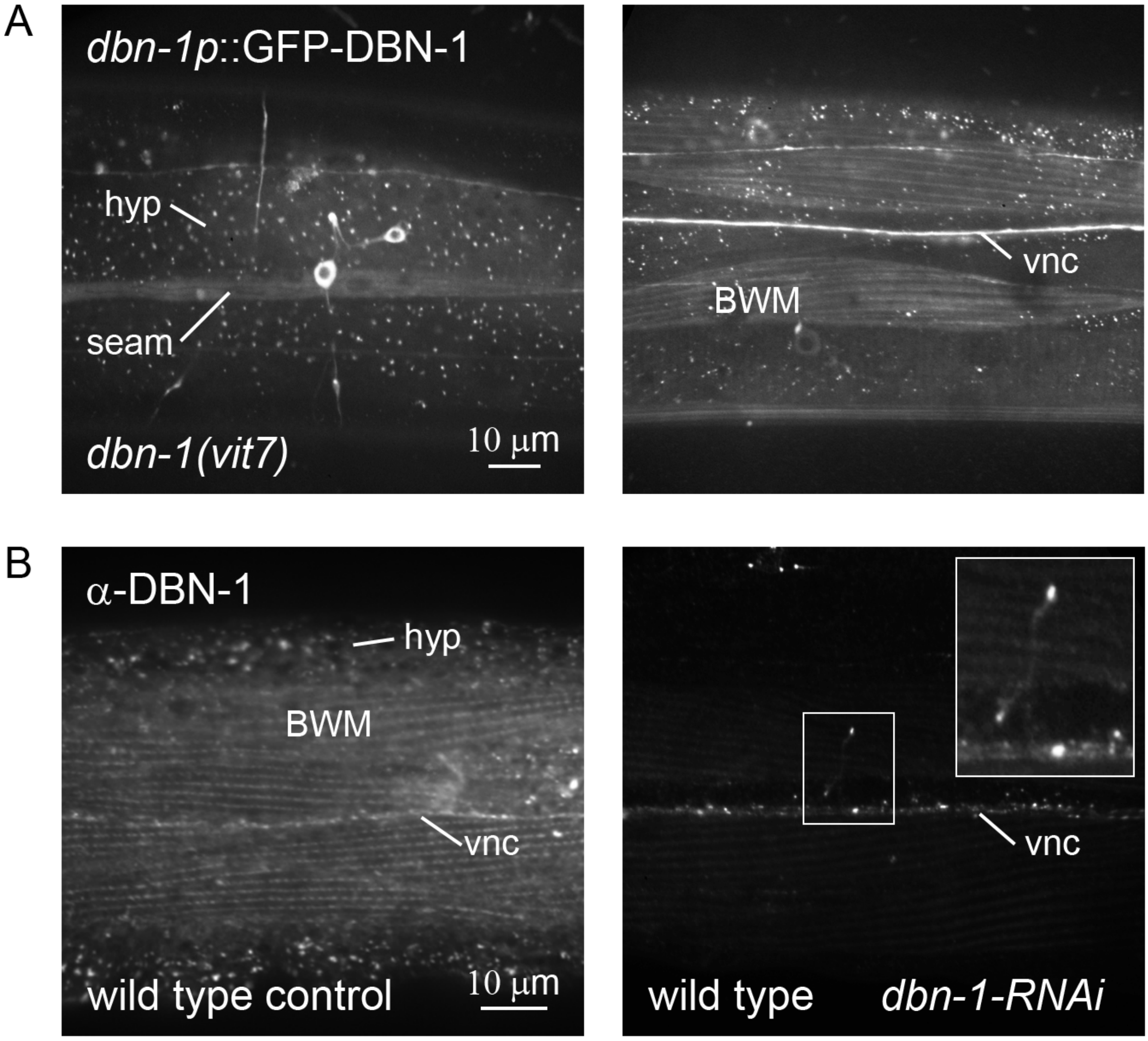


**Figure S1.** DBN-1 is expressed in multiple tissues, including body-wall muscles and neurons. **A.** Expression pattern of *dbn-1p*::GFP-DBN-1 in *dbn-1(vit7)* mutant. A GFP signal is visible in the hypodermis (hyp, punctate pattern) including seam cells, in neurons (ventral nerve cord (vnc)) and in body-wall muscles (BWM). **B.** Expression pattern of endogenous DBN-1. Wild type worms fed with control RNAi (left) or RNAi against *dbn-1* (right) and immunostained with antibodies against DBN-1. In the control sample, DBN-1 is localized in the hypodermis (hyp, punctate pattern), body-wall muscles (BWM), and the ventral nerve cord (vnc). Neuronal expression of DBN-1 remains clearly visible, while the protein is not visible in the hypodermis and in muscles after *dbn-1*-RNAi feeding.

### Supplemental methods

#### *RNA interference*

Experiments were performed as described in detail in (Butkevich et al. 2015). Briefly, young adult *C. elegans* nematodes were allowed to lay eggs overnight on NGM agar plates, coated with *E. coli* HT115 (DE3) expressing the 1-500 bp cDNA fragment of DBN-1 in L4440 plasmids. A control RNAi experiment was performed using the unmodified L4440 plasmid. F1 were then cultured at 20°C and immunostained at the young adult stage.

#### *Immunostaining*

Whole worms were fixed and permeabilized following the Nonet method (Nonet et al. 1993). Affinity purified rabbit anti-DBN-1 antibody (Butkevich et al. 2015) was used in combination with Alexa Fluor 488 goat anti-rabbit IgG (A11008; ThermoFisher Scientific).

#### *Fluorescence microscopy*

Fluorescence images were obtained using an Axiovert 200M microscope equipped with a 63x / NA 1.4 objective (Zeiss), a CSU10 spinning disk confocal imager (Yokagawa Electric Corporation), and an iXon EMCCD camera (Andor Technology).

### References:

Butkevich E, Bodensiek K, Fakhri N, von Roden K, Schaap IA, Majoul I, Schmidt CF, Klopfenstein DR. 2015. Drebrin-like protein DBN-1 is a sarcomere component that stabilizes actin filaments during muscle contraction. *Nature communications* **6**: 7523.

Nonet ML, Grundahl K, Meyer BJ, Rand JB. 1993. Synaptic function is impaired but not eliminated in C. elegans mutants lacking synaptotagmin. *Cell* **73**: 1291-1305.
